# Proline substitutions in the ASIC1 β11-12 linker slow desensitization

**DOI:** 10.1101/2024.05.09.593312

**Authors:** Rutambhara Purohit, Tyler Couch, David M. MacLean

**Affiliations:** Department of Pharmacology and Physiology, University of Rochester Medical Center

**Keywords:** Acid-sensing ion channels, ASICs, gating, desensitization, concatemers

## Abstract

Desensitization is a prominent feature of nearly all ligand gated ion channels. Acid-sensing ion channels (ASIC) undergo desensitization within hundreds of milliseconds to seconds upon continual extracellular acidification. The ASIC mechanism of desensitization is primarily due to the isomerization or “flipping” of a short linker joining the 11^th^ and 12^th^ beta sheets in the extracellular domain. In the resting and active states this β11-12 linker adopts an “upward” conformation while in the desensitized conformation the linker assumes a “downward” state. To accommodate this “downward” state, specific peptide bonds within the linker adopt either trans-like or cis-like conformations. Since proline-containing peptide bonds undergo cis-trans isomerization very slowly, we hypothesized that introducing proline residues in the linker may slow or even abolish ASIC desensitization, potentially providing a valuable research tools. Proline substitutions in the chicken ASIC1 β11-12 linker (L414P and Y416P) slowed desensitization decays approximately 100 to 1000-fold as measured in excised patches. Both L414P and Y416P shifted the steady state desensitization curves to more acidic pHs while activation curves and ion selectivity of these slow-desensitizing currents were largely unaffected. To investigate the functional stoichiometry of desensitization in the trimeric ASIC, we created families of L414P and Y416P concatemers with zero, one, two or three proline substitutions in all possible configurations. Introducing one or two L414P or Y416P mutations only slightly attenuated desensitization, suggesting that conformational changes in the remaining faster wild type subunits were sufficient to desensitize the channel. These data highlight the unusual cis-trans isomerization mechanism of ASIC desensitization and support a model where a single subunit is sufficient to desensitize the entire channel.

## Introduction

Desensitization is a common property of neurotransmitter-gated ion channels (NGICs)^1^ where continual exposure to agonist terminates ion flow. It is thought to be a protective mechanism, curtailing ion influx when transmitter release fails to terminate. Consistent with this idea, eliminating channel desensitization can have adverse developmental effects in mice^2^. The structural basis for channel desensitization is becoming clearer for many NGICs, thanks to structural and functional efforts^3–8^.

Acid-sensing ion channels (ASICs) also act as NGICs with protons serving as the transmitter^9–12^. As with other NGICs, ASICs exhibit fast activation and desensitization in response to rapid extracellular pH decreases^13,14^. The desensitization time course of ASICs ranges from hundreds of milliseconds to seconds, depending on the subunit, species variant and activating pH^14,15^. Structural and functional studies indicate ASIC desensitization arises from a “molecular clutch” mechanism with a short linker connecting the 11^th^ and 12^th^ β-sheets serving as the clutch^15–18^. In the resting and toxin-stabilized open states, the β11-12 linker adopts an “upward” conformation (Figure 1A), coupling conformational changes in the upper extracellular domain to pore opening. However, in the desensitized structures this linker adopts a “downward” conformation (Figure 1A), uncoupling conformational changes and thereby allowing the extracellular domain to remain in the active protonated form while the pore closes^15,16,18^. This structural proposal has been supported by multiple lines of evidence including molecular dynamics, extensive mutagenesis, state-dependent non-canonical amino acid photocrosslinking and voltage clamp fluorometry^16–19^. While other channel regions can impact the desensitization process^20–22^, accumulating evidence points to the β11-12 linker as the critical substrate for ASIC desensitization.

**Figure 1.**
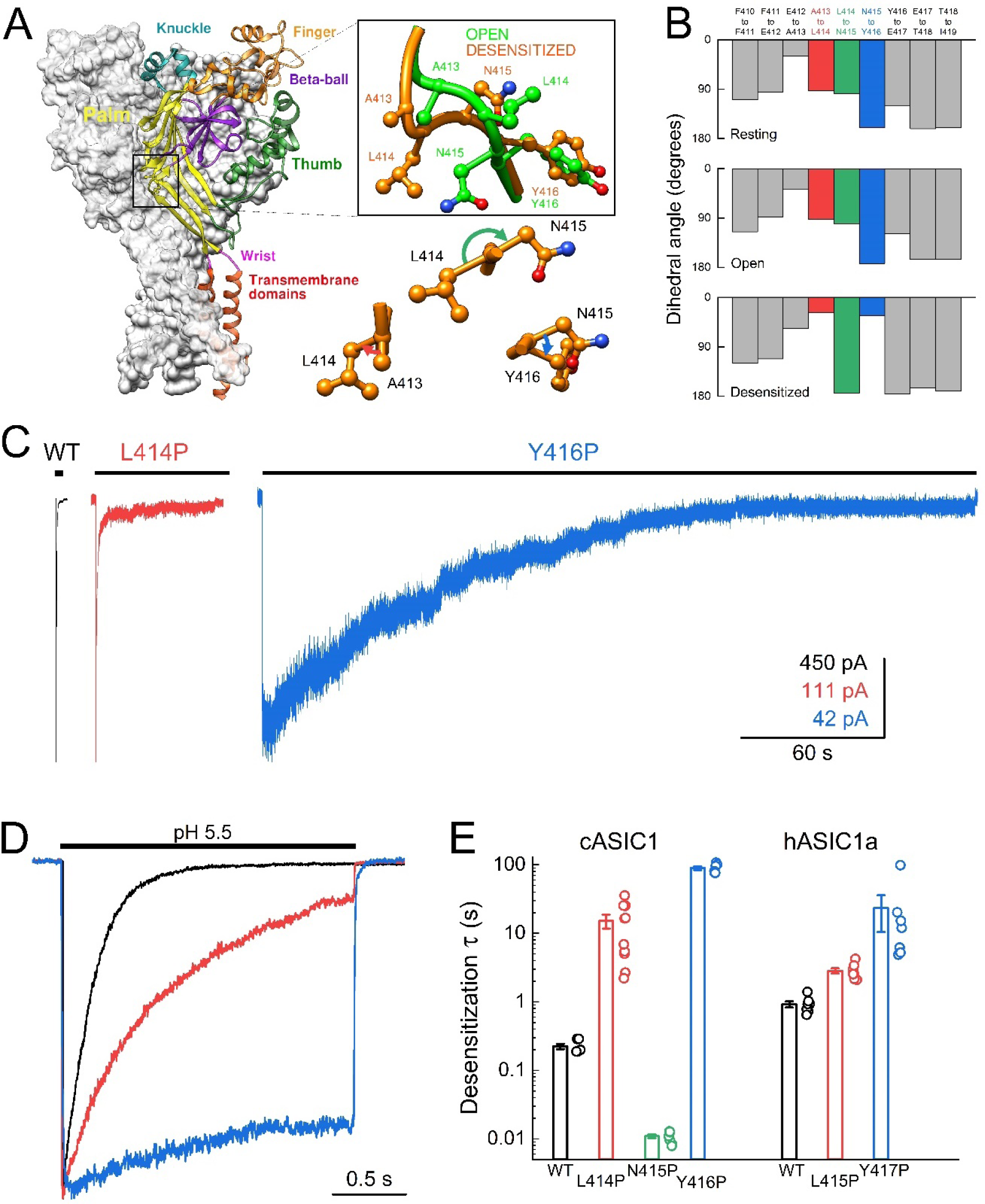
Proline substitutions into β11-12 linker can considerably slow desensitization. **(A,** *left*) Structure of cASIC1 resting state trimer (PDB: 6VTL). Domains are identified by color in one subunit while remaining two subunits are colored grey. **(A, *right*)** Boxed region shows close-up view of the β11-12 linker (boxed area in panel A, (*left*)) in the open and desensitized states (PDB: 4NTW and 6NTK). Lower panel depicts dihedral angles of β11-12 linker peptide bonds. **(B)** Dihedral angle for indicated peptide bonds in resting, open and desensitized structures (PDBs: 6VTL, 4NTW, 4VTK). **(C)** Peak normalized outside-out patch responses from cASIC1 wild-type (*black trace*), L414P (*red trace*) or Y416P (*blue trace*) channels during a 300 second jump from pH 8 to pH 5.5. **(D)** Same constructs as in C but using a 2 second jump. **(E)** Summary of weighted desensitization time constants for the indicated mutations in either cASIC1 or hASIC1a. Desensitization time constant of cASIC1: WT 0.22 ± 0.02 s, n = 6; L414P 15 ± 3 s, n = 11, p = 0.006 versus WT; N415P 0.01 ± 0.007, n = 8, p < 1e^-6^ versus WT; Y416P 89 ± 6 s, n = 6, p < 1e^-6^ versus WT. Desensitization time constant of hASIC1a: WT 0.92 ± 0.09, n = 7; L415P 2.8 ± 0.2, n = 8, p < 1e^-6^ versus WT; Y417P 23 ± 13, n = 7, p < 1e^-6^ versus WT. Circles denote individual patches and error bars show S.E.M.

In the resting and open states, the β11-12 linker peptide bonds are in the trans conformation. However, in the desensitized structures two of the linker peptide bonds adopt a cis conformation (Figure 1A and B). Peptide bonds containing C terminal proline undergo cis-trans isomerization very slowly. Indeed, the slow cis-trans isomerization of proline is proposed to act as a clock or timer to regulate protein folding^23,24^. We hypothesized that introducing proline residues at linker positions undergoing cis-trans isomerization would markedly slow, and perhaps even prevent, ASIC desensitization. Such mutations maybe useful for subsequent biophysical investigations of channel gating or perhaps inquiry into the physiological role of ASIC desensitization^2^. To test this, we measured the kinetics of ASIC desensitization in chicken ASIC1 (cASIC1) and human ASIC1a (hASIC1a) with proline substitutions. We also characterized the pH dependance and cation selectivity of these mutations. Finally, we developed a novel system to rapidly and reliably generate concatenated ASICs. We use this concatemer system to incorporate single or double proline substitutions in defined positions within the trimer and assess the consequences to the desensitization time course.

## Materials and Methods

### Cell culture and transfection

Human embryonic kidney (HEK-293T) ASIC-KO cells^25^ were maintained in Minimum Essential Medium (Gibco #11095) supplemented with 10% EqualFetal Bovine Serum (Atlas Biologicals #EF-0500-A), and 1% PenStrep (Invitrogen #15140122). Cells were grown at 37°C in a humidified atmosphere with 5% CO_2_ until approximately 80% confluent after which they were sub-cultured into 35 mm dishes for transfection. Transient transfection was done with polyethyleneimine (Polysciences #23966) using an ASIC:eGFP ratio of 10:1 and PEI:DNA ratio-3:1 according to manufacturer’s instructions. For monomeric constructs the transfection medium was replaced with fresh medium after 3 hours and recordings done after 24 hours. For transfection of concatemers, a 5:1 PEI:DNA ratio was used, the media replaced after 12 hours and recordings done after 48 hours.

### DNA constructs and mutagenesis

Mutations to cASIC1 and hASIC1a (G variant^26^) in pcDNA3.1(+) were introduced with site directed mutagenesis using non-overlapping primers. PCR was performed using Q5 HotStart high fidelity master mix (New England Biolabs #M0494S). Following KLD enzyme treatment (NEB #M0554S), XL-10 Gold ultracompetent cells (Agilent) were transformed with 5 µL of reaction mix. Colonies were screened by Sanger sequencing. Concatemers were generated with a single pot reaction combining three holding vectors, one for each subunit, with a PCR-opened destination vector, Type IIS restriction enzyme BbsI and T4 ligase with alternating cut/ligate cycles driven by temperature. To create the three holding vectors the coding sequence of cASIC1 was PCR amplified using 3 distinct primer sets with overhangs encoding unique linker sequences, BbsI sites, and NheI and XhoI sites on the forward and reverse primers, respectively. Products were cloned into pEGFP-C1 at the NheI and XhoI sites. Point mutations were introduced into these holding vectors as described above. To create the destination vector, pcDNA3.1(+) was domesticated by removing existing BbsI sites and the SV40-driven neomycin resistance gene was replaced with mNeonGreen^27^. This plasmid, termed pcDNA3.1.2(+) was PCR-opened with primers encoding BbsI sites, Dpn1 treated, and column purified. Golden Gate reactions were set up to contain PCR-derived domesticated pcDNA3.1.2(+) and each cASIC1 subunit holding vector at a 1:2:2:2 molar ratio, 10 mM dithiothreitol, 1 mM ATP, 20 units of BbsI-HF, and 400 cohesive end units of T4 DNA ligase. Reactions were thermocycled 30-45 times between 37 and 16 °C for 1 minute followed by 60 °C for 5 minutes and held at 12 °C. Cells were transformed into *E. coli Stbl3* cells with either 30 or 37 °C recovery before plating on LB agar plates with 100 µg/ml ampicillin. Plates were incubated overnight at 30°C to minimize recombination. Single colonies were expanded, miniprepped, and test digested with BamHI and BglII. All clones were confirmed using whole plasmid nanopore sequencing (Eurofins Genomics or Plasmidsaurus).

### Western Blots

The expression of monomeric and concatemeric channels was confirmed by Western blotting. Twenty-four to forty-eight hours post transfection, cells were washed with PBS then rocked in 2x Laemmli buffer (BioRad #1610737) with 5% β-mercaptoethanol for 30 minutes at room temperature. Lysates were cleared by centrifugation and then stored in −20°C. Equal volumes of protein samples were resolved on 4-12% gradient SDS-polyacrylamide gel and transferred onto 0.2μM nitrocellulose (BioRad #1620112) or PVDF (BioRad #1620177) membrane. Following blocking with 5% bovine serum albumin in TBST, blots were incubated overnight at 4°C with anti-ASIC1 primary antibody (BioLegend #833501) at a 1:1000 dilution. Following washes with TBST, membranes were treated for 1 hour at room temperature with HRP-conjugated secondary antibody (Azure #AC2115) at a 1:5000 dilution. HRP-conjugated anti-GAPDH primary antibody (BioLegend #649203) with a dilution of 1:1000 was used as an internal control. Blots were imaged in Azure Imager(c300) using chemiluminescent reaction mix (Thermo #34580).

### Electrophysiology

Outside-out patches were excised from transfected HEK-293T ASIC KO cells^25^ using heat-polished, thick-walled borosilicate glass pipettes of 3 to 6 MΩ resistance. The pipette internal solution contained (in mM) 135 CsF, 33 CsOH, 11 EGTA, 10 HEPES, 2 MgCl_2_ and 1 CaCl_2_ (pH 7.4). The external solution was composed of (in mM)150 NaCl, 10 HEPES (or MES for solutions of final pH value less than 7), 1 CaCl_2_ and 1 MgCl_2_. The pH was adjusted using Tris-base. Unless otherwise stated, external solutions of pH 8.0 and pH 5.5 were used as baseline and stimulus solutions, respectively. All recordings were performed at room temperature with a holding potential of −60 mV using an Axopatch 200B amplifier and Clampex software (Molecular Devices) with a sampling frequency of 20-50 kHz, 10 kHz filter and digitized using 1550 (Molecular Devices). Series resistance was routinely compensated by 90 to 95% when the peak amplitude exceeded 100 pA. Rapid perfusion was performed using double or triple-barrel application pipettes (Vitrocom)^28^ with E-421 piezo-driven translators (Physike Instruments). Piezo voltage commands were low-pass filtered at 30-150 Hz (8-pole Bessel, Frequency Devices). Unless otherwise stated, at least 10 seconds (and often greater) of pH 8 solution separated successive acidic pH stimuli. For concentration response curves, responses to pH 5.5 were interleaved with responses to sub-saturating stimuli to control for rundown. Similarly, for steady-state desensitization experiments (SSD, Figure 2), responses from conditioning pH 8 were interweaved with responses from more acidic conditioning pHs. Patches were incubated in conditioning pHs for 60 seconds before application of stimulating pH. For bi-ionic reversal potential measurements (Figure 1 – figure supplement 1), the internal solution composition was (in mM) 135 NaCl, 11 EGTA, 10 HEPES, 2 MgCl_2_ and 1 CaCl_2_. The external solution was either the standard above or had the NaCl replaced with equimolar KCl. Acid activated currents were measured with holding potentials of −100 mV to + 80 mV with 20 mV increments.

**Figure 2.**
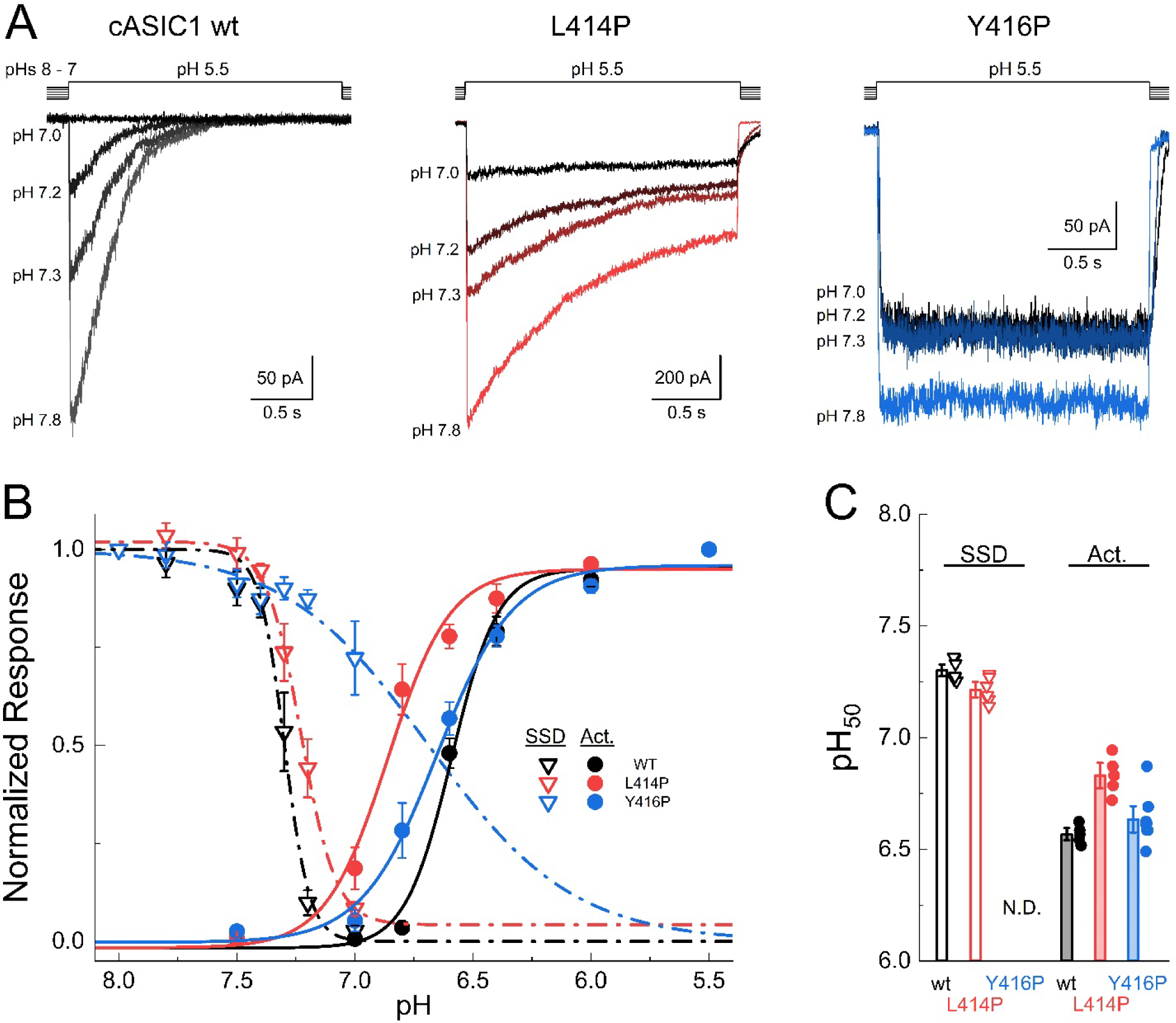
Characterization of L414P and Y416P pH-dependence. **(A)** Outside-out patch responses of wild type (*left, black*), L414P (*middle, red*) and Y416P (*right, blue*) to pH 5.5 after pre-incubating with conditioning pH ranging from pH 8.0 to pH 7.0. **(B)** Summary of pH response curves for SSD (*solid lines*) and activation (*dashed lines*) for wild type (*black*), L414P (*red*) and Y416P (*blue*). Symbols show mean values across patches. **(C)** Summary of pH_50_s of SSD and activation of all patches of wild type, L414P and Y416P. SSD pH_50_s: WT 7.3 ± 0.02, n = 7; L414P 7.2 ± 0.02, n = 6, p = 0.016 versus WT. Activation pH_50_s: WT 6.56 ± 0.01, n = 5; L414P 6.80 ± 0.03, n = 5, p < 1e^-6^ versus WT; Y416P 6.61 ± 0.01, n = 8, p = 0.24 versus WT. Symbols represent individual patch values. All error bars are SEM. N.D. stands for not determined.

### Statistics and Data Analysis

Current desensitization decays were fitted using single or double exponential decay functions in Clampfit (Molecular Devices). Here we report the weighted time constant to enable easier comparison. For concentration response curves, peak currents in response to the activating pH were normalized to the nearest peak current evoked by pH 5.5. Similarly, for SSD experiments the responses obtained with various conditioning pHs were normalized to those from the nearest conditioning pH 8 response. All patches were fit individually with the following function:

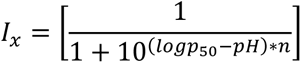

Here ‘I_x_’ is the normalized current response at a particular ‘pH’, ‘pH_50_’ is the pH at half-maximal response and ‘n’ is the slope. The mean pH_50_s were used to evaluate statistical significance. The permeabilities of external [Na^+^] or [K^+^] relative to internal [Na^+^] (P_x_/P_Na_) was calculated using the following function-

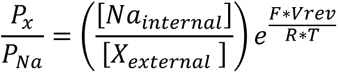

Here, Na_internal_ and X_external_ are concentrations of internal [Na^+^] and external [Na^+^] or [K^+^]. ‘V_rev_^’^ is reversal potential in volts and F, R and T have their usual thermodynamic meanings.

All column bars show mean ± S.E.M. ‘n’ represents the number of patches. Unless otherwise noted, statistical testing was done using nonparametric two-tailed, unpaired randomization tests with at least 100,000 iterations implemented in Python to assess statistical significance. Concatemer kinetics and concentration response curves were assessed in Prism using 1-way Welch’s ANOVA with Dunnett’s post-hoc multiple comparisons or 2-way ANOVAs (assuming no interaction effect) with Holm-Sidak post-hoc multiple comparisons.

## Results

### Proline substitution at Leu414 and Tyr416 slows desensitization

ASICs assemble as homo or heterotrimers with each subunit containing a large ∼300 amino acid extracellular domain flanked by single transmembrane helices and intracellular N and C terminal tails. The extracellular domain resembles a hand, consisting of wrist, palm, knuckle, finger and thumb domains as well as a β-ball domain (Figure 1A)^16,29^. The palm domain can be further separated into an upper and lower region with the β11-12 linker in between these two regions (Figure 1A, boxed region). In the resting and open states, the side chains of β11-12 linker residues Leu414 and Asn415 point upwards and downwards, respectively (Figure 1A, cASIC1 numbering)^16,30^. In the desensitized state, both Leu and Asn side chains essentially swap their positions with the Leu side chain pointing downward and the Asn pointed upwards (Figure 1A)^31^. Interestingly, at some positions within the linker the side chain reorientation appears to be a cis-trans isomerization. To illustrate this, we measured the dihedral angle between side chains (defined by C_β_-C_α_ to C_α_ -C_β_) through the β11-12 linker in resting, toxin-stabilized open and desensitized states (PDB: 6VTL^32^, 4NTW^30^, 6VTK^32^, respectively). In both resting and toxin-stabilized open states, all but one dihedral angles is approximately 90° or greater, consistent with a more trans conformation (Figure 1B). However, in the desensitized state the A413 - L414 and N415 - Y416 dihedral angles dropped to near 0°, indicative of a cis conformation (Figure 1A and B, red and blue). Such movements should principally arise from rotation of either phi or psi bonds since the peptide bond itself has pseudo-double bond character. Thus, restricting phi or psi bond rotation should slow linker isomerization. Proline has an extremely rigid phi bond owning to the pyrrolidine ring. Therefore, we reasoned that substituting proline into various linker positions should slow isomerization and consequently slow or even eliminate channel desensitization.

To test this, we measured desensitization kinetics of cASIC1 wild type, L414P, N415P and Y416P in outside-out patches with jumps from a conditioning solution of pH 8 into a maximally activating solution of pH 5.5 (Figure 1C and D). Consistent with our hypothesis, proline substitution at both L414P and Y416P desensitized far more slowly than wild type channels, nearly 100x slower for L414P and nearly 1000x fold slower in the case of Y416P (Figure 1C-E). The considerable slowing of desensitization required using much longer 300 second agonist jumps to properly measure kinetics but was still visible on shorter 2 second jumps (Figure 1C and D). Proline substitution into hASIC1a at the analogous positions produced similar slowing of desensitization (Figure 1E), however the effect was smaller than that observed in cASIC1. Interestingly, cASIC1 N415P actually accelerated the desensitization time course (Figure 1E). Unfortunately, we could not detect currents from hASIC1a N415P despite repeated attempts. Our interpretation of the faster kinetics of cASIC1 N415P is that since the L414 to N415 peptide bond remains in trans for both resting and desensitized states then restricted phi movement of N415P essentially links the movements of L414 and N415P together (more in ‘Discussion’).

### Cation selectivity is unchanged in proline mutants

Normal acid-evoked cASIC1 currents are Na^+^ selective^13,33,34^. However, currents evoked by more mild acidic stimuli and in the presence of PcTx1 are non-selective^35^. The mild acid/PcTx1 evoked currents are also slower than normal currents^35^ and resemble that of L414P and Y416P responses. Given the capacity for cASIC1 to lose cation selectivity, we measured the relative selectivity of wild type, L414P and Y416P mutations using bi-ionic reversal potential measurements (Figure 1 – figure supplement 1). Using a Na^+^ based internal solution, currents evoked by pH 5.5 solutions were measured with external Na^+^ or K^+^ over a range of voltages (−100 mV to + 80 mV, 20 mV steps) in the same patch (Figure 1 – figure supplement 1)^33^. The reversal potentials in either external Na^+^ or K^+^ were indistinguishable across mutations (Figure 1 – figure supplement 1B). The relative permeabilities of K^+^ or Na^+^ with respect to Na^+^ were also not significantly different (Figure 1 – figure supplement 1C). Therefore, slowing desensitization via proline substitutions in the β11-12 linker does not alter cation selectivity.

### L414P and Y416P slightly alter activation and steady state desensitization

To determine if β11-12 linker proline substitutions altered the pH dependance of activation, we constructed pH response curves using a conditioning solution with a pH of 8 and activating solutions with pH’s ranging from 7.5 to 5.5. Consistent with past work, cASIC1 wild type had an activation pH_50_ of approximately 6.5 (pH_50_ 6.56 ± 0.01, n = 5, Figure 2). The Y416P mutation did not markedly alter the pH50 of activation (6.61 ± 0.01, n = 8, p = 0.24 versus WT), interestingly however the L414P mutation left shifted the activation pH_50_ by 0.24 pH units (6.80 ± 0.03, n = 5, p < 1e^-6^ versus WT). Next, we examined the pH_50_ of steady-state desensitization (SSD). Patches were incubated in extracellular solutions with pH values ranging from 8 to 7 for 60 seconds before a test pulse of pH 5.5 (Figure 2A). Consistent with past data, the SSD pH_50_ of wild type was 7.3 ± 0.02 (n = 7). Since L414P and Y416P slow desensitization evoked by pH 5.5 applications by orders of magnitude (Figure 1), these mutations likely also slow entry into SSD by a comparable amount. Thus, properly measuring equilibrium SSD requires lengthening the conditioning pulses by orders of magnitude. This is not experimentally feasible, particularly across multiple conditioning pHs. Instead, we chose to compare SSD at a common conditioning time of 60 seconds, with the caveat that these SSD pH_50_s are not necessarily at equilibrium. Under these conditions, L414P right shifted the SSD pH_50_ by 0.1 pH units to 7.2 ± 0.02 (n = 6, p = 0.016). Notably, SSD of Y416P was minimal even with conditioning pH solutions as acidic as 7 (Figure 2A and B). Further, the minimal SSD of Y416P precluded accurate pH50 determinations for each patch (Figure 2C). In summary, the pH dependance of activation is only slightly altered by the L414P mutation. And, as expected of mutations that make desensitization less favorable, the SSD curves for L414P and Y416P are right shifted or the onset of desensitization at these mild acidities is so slow it’s not feasible to measure.

### Contribution of each proline linker to delayed desensitization

The prominent slowing of desensitization by proline substitutions provided a tool to assess the contribution of individual subunits to desensitization. By progressively introducing one, two or three proline mutations into trimeric ASICs and assessing the relative changes in desensitization, we can address how many subunits must adopt a desensitized conformation for the entire channel to inactivate. We turned to concatemeric ASICs as a means of introducing a defined number of mutations. Concatemeric ASICs have been used in the past^36–40^, however their assembly can be challenging. We sought to develop a flexible and reliable concatemer system that could be used for the present purpose and beyond. To that end, we constructed three ‘holding vectors’, each containing a single cASIC1 subunit termed Subunit A, B or C. The coding region of each subunit is flanked by BbsI sites, a Type IIS restriction enzyme that cleaves outside its recognition site (Figure 3A). Digesting these holding vectors with BbsI exposes unique four basepair overhangs that complement the adjacent subunits. We combined these holding vectors with a PCR-derived backbone containing BbsI 5’ and 3’ sites into a single pot Golden Gate reaction. Alternating between 37°C, to drive BbsI activity, and 16°C, to drive T4 ligase, results in an assembled concatemer expression plasmid with Subunits A, B and C in order, no BbsI sites and antibiotic resistance of the PCR-derived backbone, not the holding vectors (Figure 3A). Proper assembly can be easily screened by double restriction digest using BamHI and KpnI followed by whole plasmid sequencing of positive clones. The concatenated cASIC1 wild type (3xWT) behaved comparably to monomeric wild type, with indistinguishable desensitization kinetics (τ_Desen_ WT: 220 ± 20, n = 6; 3xWT: 210 ± 15, n = 8, p = 0.7) and comparable pH_50_s of SSD and activation (although a 0.1 pH unit right shift in the SSD pH_50_ was detected in the 3xWT, Figure 3B - D). However, as expected, the peak currents from the 3xWT were smaller than that of monomeric wild type cASIC1 (3xWT peak current: 1.2 ± 0.3 nA, n = 10; WT: 2.8 ± 0.4 nA, n = 8, p = 0.014). These observations indicate that linking of subunits in the intracellular side does not markedly affect opening and desensitization kinetics of the channel although the functional expression is decreased, either due to reduced transfection efficiency of the larger plasmid, impaired biogenesis and trafficking or both.

**Figure 3.**
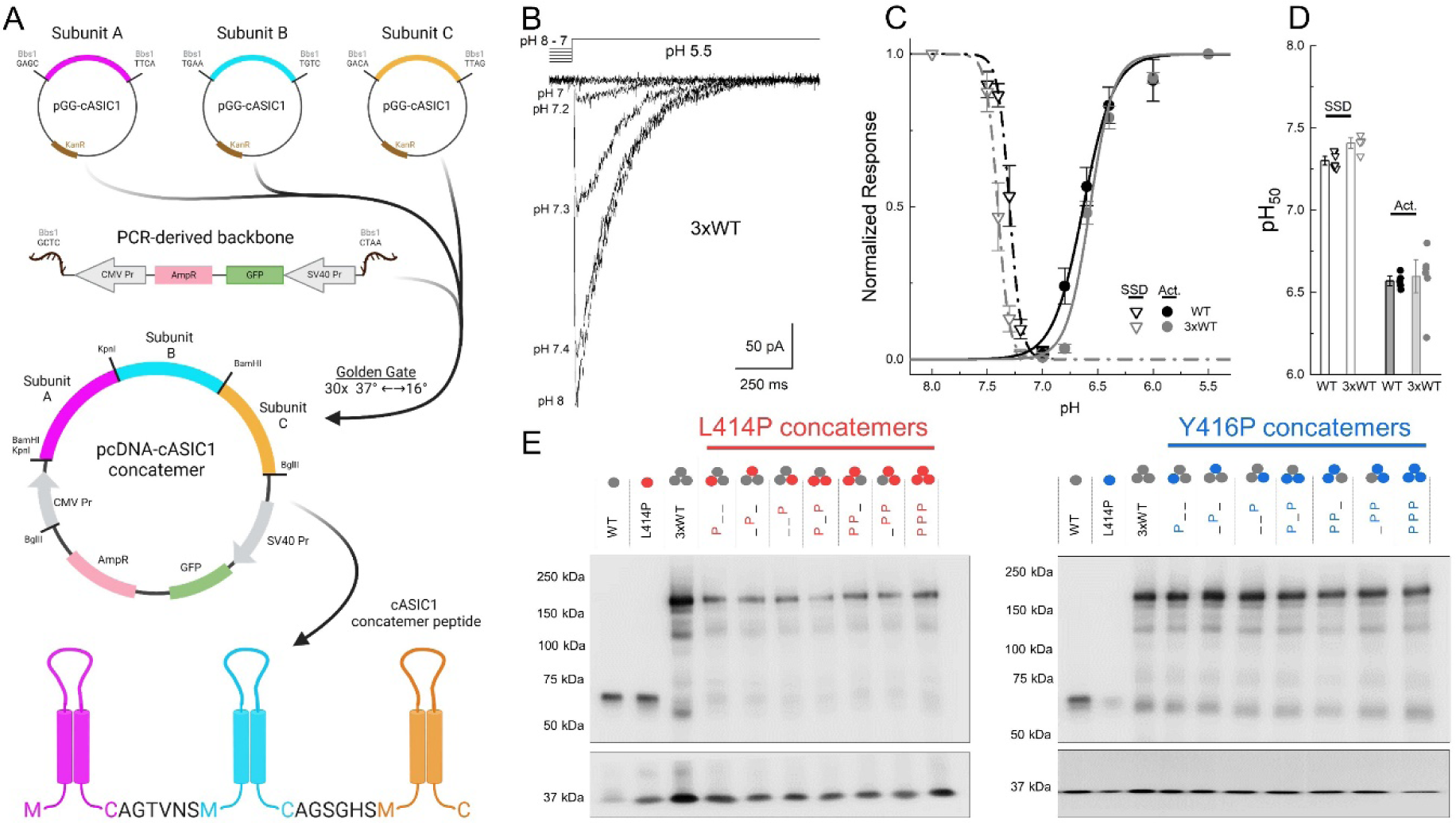
Construction and characterization of cASIC1 concatemers. **(A)** Schematic of concatemer system. Three kanamycin resistant holding vectors contain cASIC1 subunits flanked by BbsI restriction sites (*upper panel*). Combining holding vectors with PCR-derived backbone template in a Golden Gate reaction yields a full ampicillin resistant plasmid with high efficiency (*middle panel*). The resulting trimeric cASIC1 contains unique linkers between subunits (*lower panel*). **(B)** Example traces from SSD experiments with cASIC1 wild type concatemers (3xWT). **(C)** pH response curves for wild type monomers (*black symbols and lines*) and concatemers (*grey symbols and lines*) for both SSD (*inverted triangles*) and activation (*circles*). Symbols are averages across patches. **(D)** Summary of pH50s for SSD and activation curves shown in **C**. SSD pH_50_ WT: 7.30 ± 0.02, n=7; 3xWT: 7.4 ± 0.02, n=5, p = 0.008. Activation pH_50_ WT: 6.56±0.02, n=5; 3xWT: 6.59 ± 0.06, n=7, p = 0.77. Symbols are individual patches and error bars are S.E.M. **(E)** Blots for cells transfected with the indicated construct. The upper blots are for ASIC and the lower blots are GAPDH loading controls. Note the most abundant band is consistent with a trimer, suggesting the majority of the population is concatenated trimers.

To determine if concatenated ASICs were stable at the protein level, we performed western blots with whole cell lysate samples from monomeric wild type, L414P and Y416P transfected cells as well as concatemers of zero, one, two or three proline substitutions at all three positions (Figure 4E). We did observe some faint bands consistent with monomers or dimers in all concatemer constructs. However, the predominant band appeared above 150 kDa consistent with the calculated molecular weight of 180 kDa of trimeric ASIC1 (Figure 4E). Thus, our 3xWT concatemer behaved as expected and all concatemers with one, two or three prolines at either L414 or Y416 also expressed, as measured in western blots. We next considered various models of ASIC desensitization and the predictions each would yield when one or two slowly desensitizing subunits is introduced.

**Figure 4.**
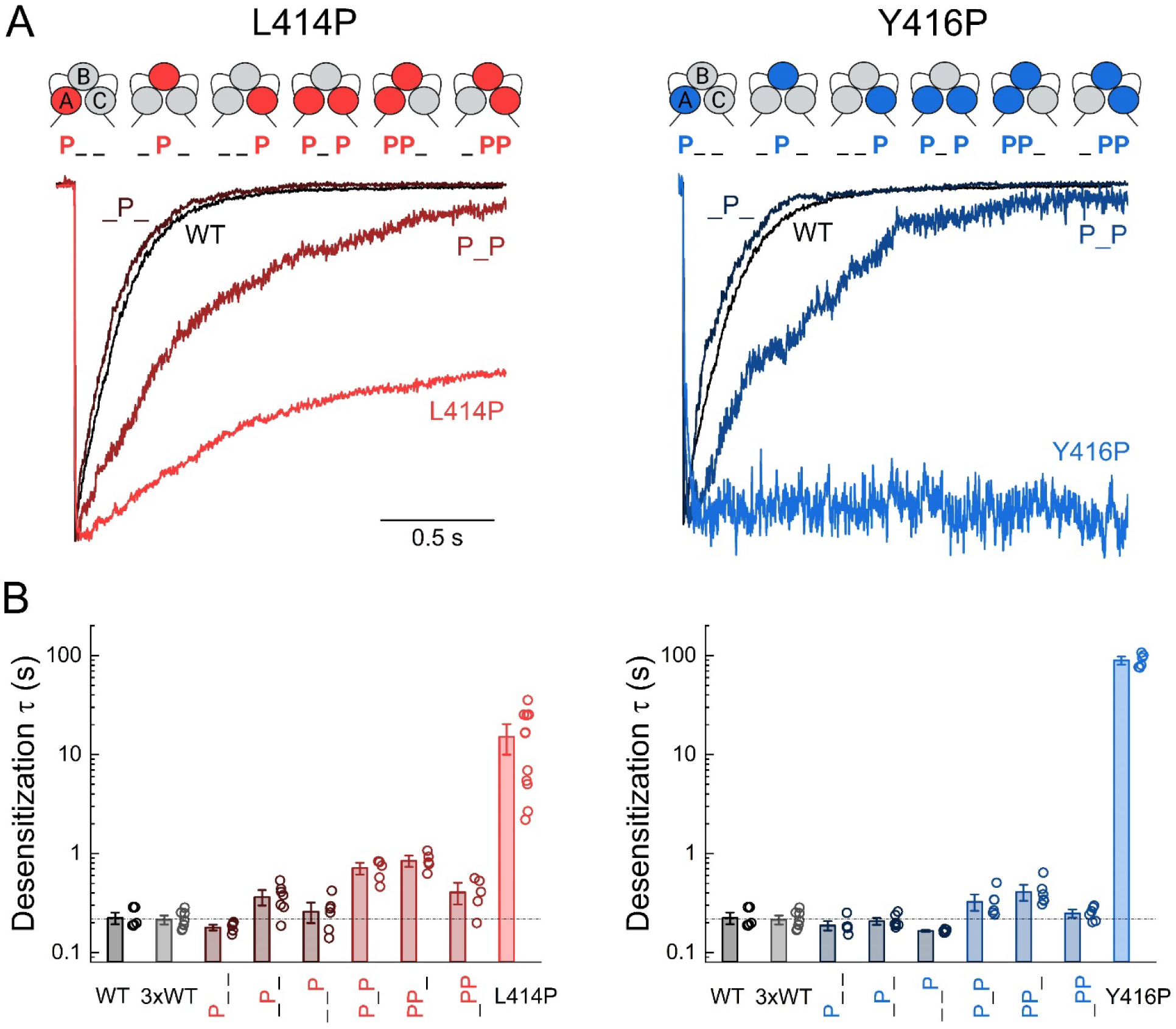
Single or double Proline substitution does not substantially slow desensitization kinetics. **(A,** *upper*) Cartoon representation of the various L414P (*left*) and Y416P (*right*) concatemers tested. **(A,** *lower)* Desensitization decays from wild type (*black*), L414P (*red*) or Y416P (*blue*) as well as the indicated Proline containing concatemer. **(B)** Summary of desensitization times constants across patches for wild type monomers (*black*) and concatemers (*grey*) as well as Proline containing concatemers and either L414P or Y416P monomers (*red* and *blue*, respectively). Symbols are individual patches, error bars are S.E.M and the dotted lines show the mean of wild type kinetics. Data for L414P and Y416P were analyzed using separate 1-way ANOVAs including 3xWT, all concatemers and the homomeric L414P or Y416P.

### Predictions of single subunit and all-subunit models of ASIC desensitization

To investigate the contribution of individual subunits to ASIC desensitization, we first considered two models of desensitization: a Single subunit model (SS) and an All-subunit model (AS). In the Single subunit model, desensitization of a single subunit is sufficient to desensitize the entire channel complex and thus desensitization kinetics will be driven by the fastest desensitizing subunit. Introducing one or two slow mutations like Y416P will produce minimal slowing of channel desensitization since the other faster wild type subunits remain to drive the desensitization reaction. In the All-subunit model, all three subunits must adopt the desensitized conformation to desensitize the channel. Desensitization of one or two subunits produces no effect on the conductance or open probability of the entire complex. Thus, desensitization kinetics will be limited by the slowest desensitizing subunit. In the AS model, introducing a single slow mutation would considerably slow desensitization as this slowest subunit is rate limiting. To provide empirical support for this reasoning, we turned to kinetic modeling.

We began with the SS model by using a branching structure where channel opening and desensitization stem from a shared fully protonated state, called R6 here (Figure 3 – figure supplement 1). The binding reaction is a multi-step co-operative binding scheme adapted from prior work^14,33^. This more complicated binding reaction is not necessary in this case (a single or two step binding reaction works just as well) but serves a larger longer-term goal of developing a robust kinetic model for ASICs. To simulate desensitization at the single subunit level, any channel entering the R6 state can then desensitize by one of three routes, representing desensitization or flipping reactions at each subunit independent of the other two (Figure 3 – figure supplement 1A). This model was fitted to wild type cASIC1 data activation curve (Figure 3 – figure supplement 1B) and desensitization decay data (Figure 3 – figure supplement 1C) and can broadly reproduce the activation pH_50_ and kinetics of desensitization (Figure 3 – figure supplement 1B and C). The same model was then fitted to Y416P desensitization data by varying only the desensitization rate, σ. We thus obtained two separate rate constants: σ_WT_, the single subunit desensitization rate constant for wild type and σ_Y416P_, the single subunit desensitization rate constant for Y416P subunits (see Table 1). We then simulated the desensitization decay of channels with two σ_WT_ subunits and one σ_Y416P_ subunit (Y2:1P) or vice versa (Y1:2P). These simulations predict that if the single subunit model is true, then introducing one or two very slow desensitizing subunits will have only a small effect on the macroscopic desensitization decay (Figure 3 – figure supplement 1C, upper). To model the ‘All-subunit’ scenario, we connected all the desensitized states except D7 to separate open states of the same conductance and rate constants as the main open state (Figure 3 – figure supplement 1A). This AS model was then re-fitted to wild type data where it better captured the slope of the activation curve (Figure 3 – figure supplement 1B). However, the AS model also resulted in slower than optimal desensitization decays during the initial phase of desensitization and faster decays in the later phase. We also fit the Y416P data and thereby obtained σ_WT_ and σ_Y416_ for this AS model (Table 1). As above, we simulated desensitization decays using one or two of each subunit. This all ‘All-subunit’ model predicts that introducing a single slow subunit is sufficient to produce substantial slowing of channel desensitization (Figure 3 – figure supplement 1C, lower, Y2:1P), with two subunits producing nearly the same effect as Y416P monomers, at least when examining the 2 second application protocols (Figure 3 – figure supplement 1C, lower, Y1:2P). For simplicity and because of the unexpectedly large variance in L414P kinetics (Figure 1E), we only modeled the desensitization of wild type and the stronger Y416P mutation but identical predictions are expected from L414P concatemers. Next, we tested these predictions by measuring desensitization decays from a range of concatemers.

**Table 1.**

|  | Single subunit model |  |  | All subunit model |  |  |
| --- | --- | --- | --- | --- | --- | --- |
|  | cASIC1 | L414P | Y416P | cASIC1 | L414P | Y416P |
| $k_{on}$ | $3 \times 10^8$ | | | | | |
| $k_{off}$ | 100 | | | | | |
| $g$ | 1.11253 | | | | | |
| $f$ | 1.41233 | | | | | |
| $\sigma$ | 9.76889 | 2.69015 | 0.170138 | 55.9006 | 14.956 | 1.75665 |
| $\gamma$ | 0.001 | | | | | |
| $\beta$ | $1 \times 10^7$ | | | | | |
| $\alpha$ | $2 \times 10^6$ | | | | | |

### Desensitization decays from L414P and Y416P concatemers

Desensitization decays to pH 5.5 evoked responses were measured for wild type, 3xWT as well as either L414P or Y416P containing concatemers with one or two proline substitutions in all possible combinations of subunits. Measuring currents from concatemers with 3 prolines was not feasible as these currents were very small (∼20-30 pA), the patches tended to be leakier and the cells had more apparent cell death. Given this, and since 3xProline concatemers should approximate monomeric Proline mutations, we focused on concatemers with only one or two prolines. Including a single copy of L414P or Y416P did not substantially alter the desensitization decays compared to wild type or 3xWT (τ_Desen_ single L414P means ± SEM range from 180 ± 7 ms to 360 ± 40 ms; Figure 4), consistent with the Single subunit model. Including two copies of L414P or Y416P did slow desensitization decays but the effect was small compared to that observed with three proline substitutions (τ_Desen_ two L414P means ± SEM range from 406 ± 60 ms to 840 ± 70 ms, Figure 4). Importantly, we observed the same pattern of effect between both L414P and Y416P in that one or two proline substitutions did not produce sizeable effects on desensitization. All three subunits must be slowly desensitizing for the entire channel to desensitize slowly (see below for details on statistical analysis). This pattern is consistent with the Single subunit model of channel desensitization where the macroscopic decay is driven mainly by the fastest subunit. However, as discussed below, we cannot rule out more complicated scenarios or models.

To statistically assess kinetic differences, we conducted 1-way ANOVAs including 3xWT data, all single and double proline concatemers for L414P (or Y416P) and the homomeric L414P (or Y416P). Data from each of these groups was normally distributed (assessed by Shapiro-Wilk) and each ANOVA was statistically significant (L414P F(7,16.93) = 21.98, p < 0.0001; Y416P F(7,16.45) = 39.85, p < 0.0001). In post-hoc testing, the homomeric Proline was different from all other groups (eg. L414P was different from 3xWT and all other L414P concatemers), supporting the Single subunit model. However, we noted that for both L414P and Y416P constructs containing a middle Proline mutation tended to have slower kinetics than other concatemers with the same number of Prolines. To further explore this apparent trend, we conducted a 2-way ANOVA on all single Proline concatemers with mutation (L414P vs Y416P) and position (P__vs__P_vs___P) as the two factors. A similar 2-way ANOVA on all double Proline concatemers was also performed. As expected, both 2-way ANOVAs detected a significant effect of having L414P versus Y416P mutations (single Prolines (F(1, 34)) = 12.47, p = 0.0012; double Prolines (F(1, 30)) = 47.38, p <0.0001)). Interestingly, both 2-way ANOVAs also detected a significant effect of position (single Prolines (F(2, 34)) = 6.66, p = 0.0036; double Prolines (F(2, 30)) = 11.96, p = 0.0002). The mechanism underlying this apparent positional effect is presently unclear (see Discussion).

To confirm that our pH 5.5 stimulus is saturating for all constructs, we constructed pH response curves. We found that all Proline containing concatemers had nearly identical pH_50_s, confirming that pH 5.5 is saturating (Figure 5). In addition, we also measured the pH_50_ of SSD for all concatemers. The pH_50_s for both SSD and activation across all concatemers were quite similar to those of wild type or 3xWT. We did find that SSD curves of Y416P concatemers were well constrained (Figure 5B), as opposed to SSD curves for Y416P homomers which could not accurately be measured (Figure 2). This is further support that single subunits are sufficient to drive appreciable desensitization.

## Discussion

Here we probed the desensitization mechanisms, and functional stoichiometry of ASIC desensitization, using proline substitutions within the β11-12 linker of cASIC1. We found that proline substitutions at L414P and Y416P produced substantial slowing of cASIC1 desensitization (Figure 1). Smaller effects were observed with hASIC1a (Figure 1). Interestingly, L414P showed a slight left shift in the activation curve while Y416P did not (Figure 2). As expected of slowly desensitizing mutations, L414P right shifted the SSD curve and Y416P desensitized so slowly SSD could not readily be measured (Figure 2). To investigate the functional stoichiometry of ASIC desensitization, we developed a novel concatemer creation system to reliably introduce Proline mutations into specific subunits within a trimer (Figure 3). Introducing one or two Prolines at either L414 or Y416 did not markedly slow desensitization. Instead, all three subunits must contain Proline to produce the expected slowing of desensitization (Figure 4), consistent with our Single subunit to desensitize kinetic model (Figure 3 – figure supplement 1). These findings highlight the unique cis-trans isomerization mechanism of the ASIC β11-12 linker in driving channel desensitization and provide evidence favoring a Single subunit desensitization model.

### Gating characteristics of linker Proline substitutions

While cASIC1 L414P and Y416P desensitized significantly more slowly than wild type, N415P accelerated the desensitization rate (Figure 1). The reason for this acceleration is unclear. One possibility is that the physical-chemical properties of Proline substituted at N415 result in fast conformational changes at that site. This region is exquisitely sensitive to small changes in amino acid side chain chemistry^17,18^. Another possibility is that linker ‘flipping’ proceeds in two (or more) steps as is observed in the limited molecular dynamics simulations performed in this region thus far^18^. It may be that inserting a rigid Proline into the linker effectively couples the multiple steps together into one more efficient movement. Further simulations, perhaps taking advantage of recent developments in constant pH molecular dynamics^41,42^, are needed to better define the molecular motions of this critical linker. We unexpectedly found the desensitization decays of L415P to be much more variable than any other mutant or wild type (Figure 1). Since all experiments were performed on a pure ASIC KO background^25^, the heterogeneity cannot arise from mixed mutant and endogenous channels. Hence, the kinetic variation appears to be intrinsic to the mutation, possibly hinting at some differences in post-translational modification that alter gating.

### Comparison with previous studies

In prior work, we have characterized an extensive set of cASIC1 β11-12 mutations^17,18^. L414P and Y416P are the slowest desensitizing mutations yet found at either position. Indeed, Y416P is the slowest desensitizing construct we have ever studied. Importantly, the slow desensitization kinetics were observed at saturating pH conditions and do not reflect a shift in pH sensitivity or other unexpected behavior at sub-saturating conditions^25^. Interestingly, we observed a blunted effect of these mutations in hASIC1a (Figure 1E, right). Since mammalian ASIC1a subunits undergo tachyphylaxis when open^43^ and cAISC1 does not^18^, this apparent blunted effect may arise more due to tachyphylaxis in the slowly desensitizing hASIC1a mutants than actual differences in subunits between species.

By introducing one or two very slowly desensitizing mutations into a concatemer, and comparing the results to kinetic simulations, we find evidence suggesting that a single subunit may be sufficient to desensitize the entire channel. In AMPA receptors, a similar ‘single subunit is sufficient’ result was obtained from careful analysis of the time course of entering the desensitized state at low agonist concentrations^44^. However, closely related kainate receptors appear to follow a more complicated tetrameric mechanism^45^. Recently, Gielen et al., examined GABA-A desensitization using a similar mutant concatemer/kinetic modelling strategy^8^. Their data was best fit with a model where two subunits must desensitize to occlude the pore. In addition, they also found that incorporating functional coupling between subunits gave overall much better fits. Here we did not incorporate functional coupling since our Single subunit model predictions matched well with the Y416P data. However, future work combining single or multiple mutations within a concatemer system may reveal some inter-subunit interactions.

### Caveats and limitations

The major limitation of our work is the possibility that the concatemers alter biogenesis, assembly or gating. Concatemers have been useful tools in exploring channel function^8,36,38,39,46^ but they do not always behave as simply as expected^47^. Past work has found that concatemers can form trans-channel interactions or breakdown at the linkers^47^, complicating interpretation. ASIC concatemers have been found to have unusual effects where concatenated dimers, trimers and tetramers all give functional responses^37^. Here we observe some apparent lower order monomers (Figure 3E) however they represent a small fraction of total channel abundance. It is possible that these monomers form the majority of the surface population in our recordings^47^. However, we do not think this likely for several reasons. First, from past work on P2X1 receptor concatemers it is the first (and second) subunit in the trimer that tends to ‘escape’ to the surface^47^ and may contribute an over-large fraction of the measured current. If this were occurring in our data, we would expect a much larger slowing of channel desensitization with concatemers containing a Proline in the first subunit (e.g. P__versus_P_or__P, Figure 4). Further, we would expect that Y416P concatemers with P__(or PP_) configurations would nearly abolish SSD as the Y416P monomer does (Figure 2). Neither of these results are observed, giving some confidence that ‘escaped’ monomers do not represent the major fraction of our observed currents. Second, while we do observe a positional effect in kinetics (Figure 4), it is Prolines at the second position that tend to give slower kinetics, not the first position as would be expected of ‘escaped’ subunits. Third, if ‘escaped’ subunits were also a persistent problem with ASIC concatemers, one would also expect a strong positional effect with concatemers made from low and high sensitivity subunits. Yet in a recent study using concatemers of low sensitivity ASIC2a and high sensitivity ASIC3, no positional effect was observed in pH response curves^38^. Nonetheless, additional investigation into the functional stoichiometry of ASIC desensitization, possibly using multiple mutations in concatemers or another strategy, is needed to further test our conclusion that a single subunit is sufficient to desensitize the channel.

We began our study with the goal of determining if Proline substitution could abolish desensitization of ASICs. If so, such mutant channels could be useful biophysical tools. Moreover, introducing a non-desensitizing ASIC1 subunit into a mouse would provide a valuable tool to investigate the physiological role of ASIC desensitization^2^. Unfortunately, we found that both L414P and Y416P (or their hASIC1a equivalents) gave relatively reduced currents compared to wild type. Thus, any future physiological studies with these mutants must distinguish effects due to slowing of desensitization versus reduced current density or biogenesis. Moreover, ASIC heteromers may represent the major population in the central nervous system and other tissues. It remains to be determined if heteromeric ASICs also follow the same functional stoichiometry as homomeric cASIC1.

## Acknowledgements

Funding for this work was provided by NIH R35GM137951 to D.M.M. We thank Dr. Matt Rook for pilot data, Lauren Bainbridge for technical assistance and members of the MacLean lab for helpful discussion.

## Author contributions

R.P. and T.C. conducted experiments and analyzed data. D.M.M conducted simulations. All authors interpreted results and contributed to writing of the manuscript.

---

**Figure 1 – figure supplement 1.**
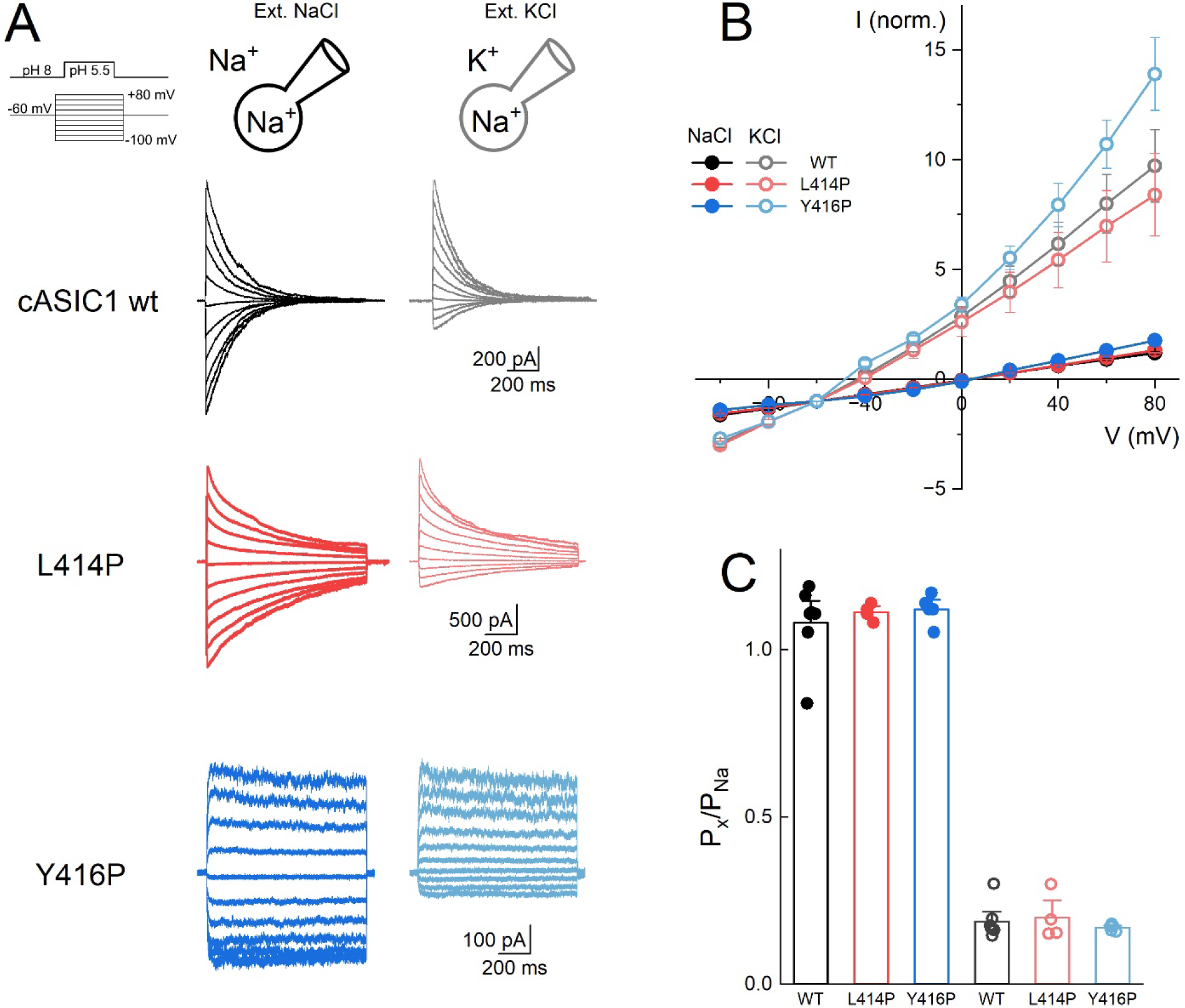
L414P and Y416P do not alter cation selectivity. **(A)** Schematic of experimental design and representative current traces of wild type (*black*), L414P (*red*) and Y416P (*blue*) during voltage steps from 100mV to +80mV in external NaCl (*left*) and KCl (*right*). **(B)** Normalized current-voltage plot of all patches of wild type (*black*), L414P (*red*) and Y416P (*blue*) in both external NaCl (*solid circles*) and external KCl (*hollow circles*). Symbols represent averages across patches. **(C)** Summary of relative permeabilities of external Na^+^ and external K^+^ with respect to internal Na^+^ for wild type, L414P and Y416P. P_Na_/P_Na:_ WT 1.08 ± 0.01, n=7; L414P 1.11 ± 0.01, n=4, p = 0.74 versus WT; Y416P 1.12 ± 0.01, n = 5, p = 0.61 versus WT. P_Ka_/P_Na_: WT 0.18 ± 0.01, n = 7; L414P 0.19 ± 0.03, n = 4, p = 0.79 versus WT; Y416P 0.16 ± 0.004, n = 5, p = 0.66 versus WT. Symbols are individual patch values. All error bars are SEM.

**Figure 3 – figure supplement 1.**
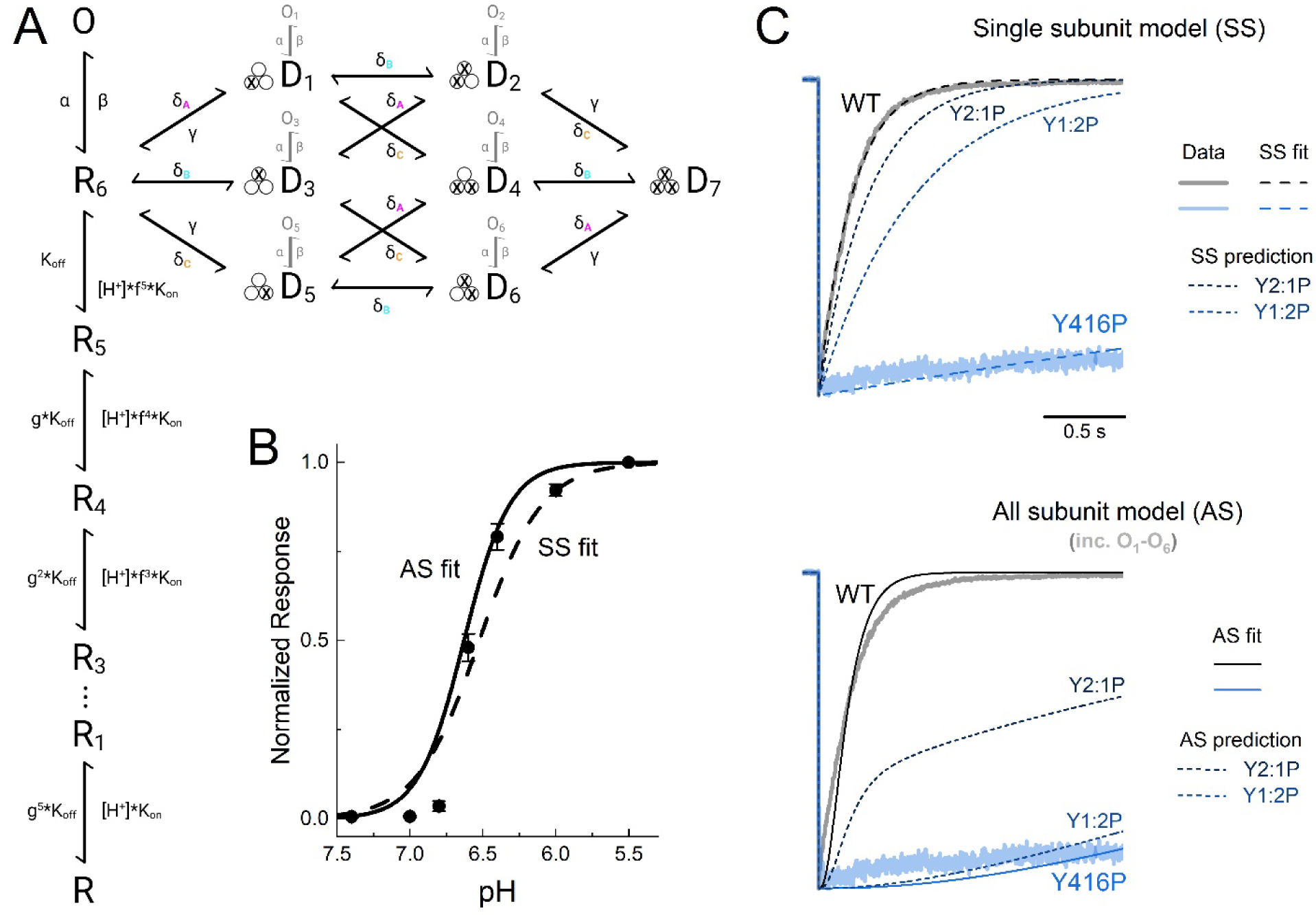
Simulations of channels with mixes of fast or slow desensitizing subunits. **(A)** Reaction mechanism with binding reaction taken from^14,33^. The Single subunit model only uses states in black. The All subunit model also includes the grey open states. **(B)** Output of Single subunit (*dashed line*) or All subunit (*solid line*) models to pH response curve for wild type cASIC1 (*black circles*). **(C)** Patch clamp data for cASIC1 wild type (*WT, transparent black*) or Y416P (*transparent blue*). Either the Single subunit model (*upper*) or All subunit model (*lower*) were fitted to this data and the output is overlayed (*dashed or solid lines, respectively*). Predictions of either Single subunit or All subunit model with the indicated number of native amino acids or Prolines are also shown (*dotted lines*).

**Supplemental Figure 3 – related to Figure 4.**
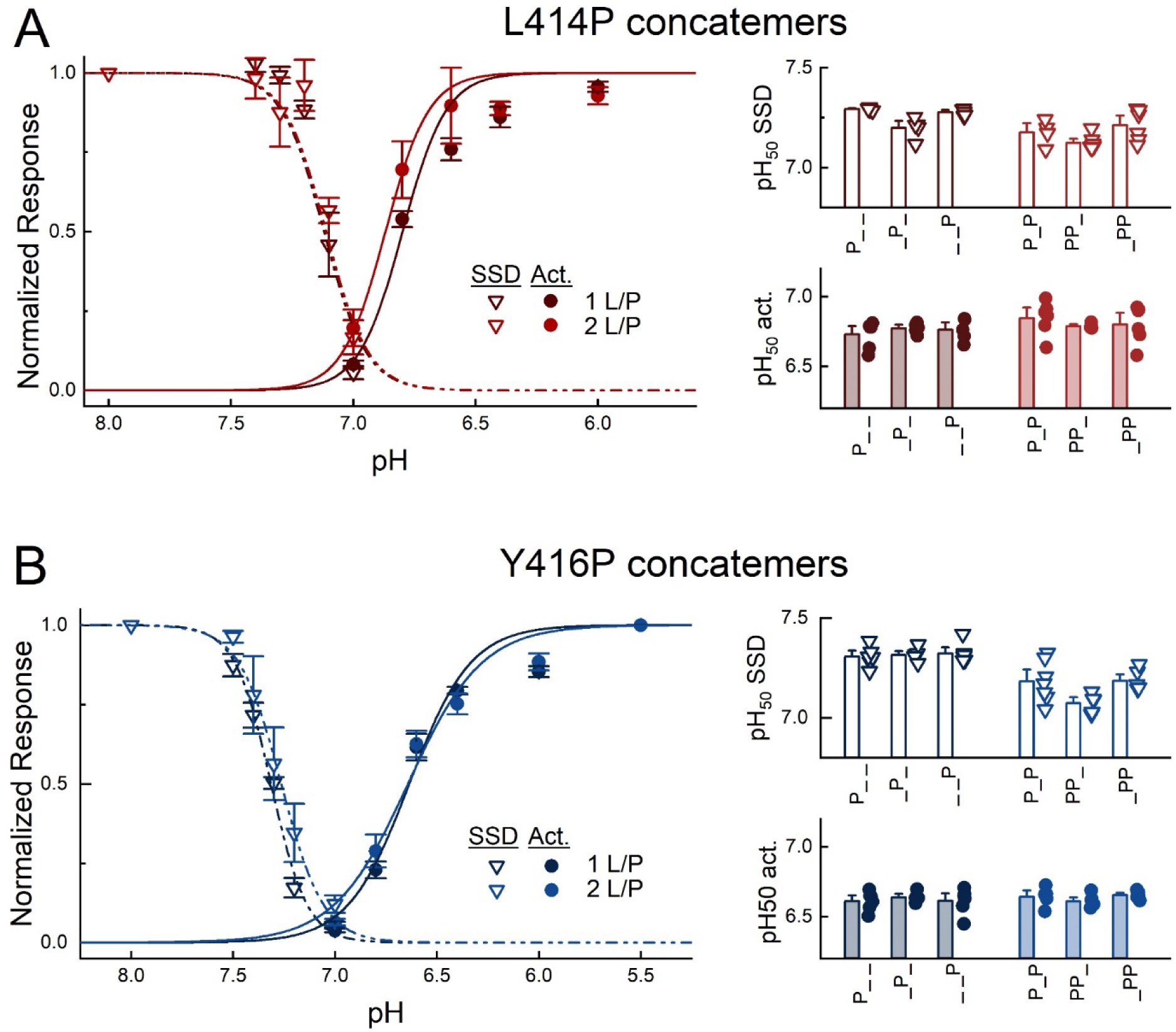
pH response curves of Proline substituted concatemers. **(A,** *left*) Response curves for SSD (inverted triangles) or activation (circles) for all L414P concatemers containing one or two Prolines. Symbols are averages across patches. **(A,** *right*) SSD and activation pH_50_s across patches for all L414P concatemers. Symbols are values for individual patches. **(B)** Same as **A** but for Y416P. All error bars are S.E.M.

